# Biological representation disentanglement of single-cell data

**DOI:** 10.1101/2023.03.05.531195

**Authors:** Zoe Piran, Niv Cohen, Yedid Hoshen, Mor Nitzan

## Abstract

Due to its internal state or external environment, a cell’s gene expression profile contains multiple signatures, simultaneously encoding information about its characteristics. Disentangling these factors of variations from single-cell data is needed to recover multiple layers of biological information and extract insight into the individual and collective behavior of cellular populations. While several recent methods were suggested for biological disentanglement, each has its limitations; they are either task-specific, cannot capture inherent nonlinear or interaction effects, cannot integrate layers of experimental data, or do not provide a general reconstruction procedure. We present *biolord*, a deep generative framework for disentangling known and unknown attributes in single-cell data. Biolord exposes the distinct effects of different biological processes or tissue structure on cellular gene expression. Based on that, biolord allows generating experimentally-inaccessible cell states by virtually shifting cells across time, space, and biological states. Specifically, we showcase accurate predictions of cellular responses to drug perturbations and generalization to predict responses to unseen drugs. Further, biolord disentangles spatial, temporal, and infection-related attributes and their associated gene expression signatures in a single-cell atlas of *Plasmodium* infection progression in the mouse liver. Biolord can handle partially labeled attributes by predicting a classification for missing labels, and hence can be used to computationally extend an infected hepatocyte population identified at a late stage of the infection to earlier stages. Biolord applies to diverse biological settings, is implemented using the scvi-tools library, and is released as open-source software at https://github.com/nitzanlab/biolord.

## Main

A cell’s gene expression profile simultaneously encodes information about multiple attributes, such as cell type, tissue of origin, and differentiation stage (Fig. 1a). Single-cell technologies can provide information about such expression profiles for cellular populations at single-cell resolution. Yet, it is still a major challenge to decode the measured gene expression, disentangling the processes from one another. A disentangled representation can uncover the existence and characteristics of diverse biological processes. It would allow the reconstruction of multiple attributes of cellular identity such as response to perturbations and infection progression. A line of earlier work suggested using factor analysis^1,2^ or non-negative matrix factorization^3^ to identify gene expression programs associated with different attributes. More recent works tackled different aspects of the disentanglement challenge for single-cell data, including computational methods that specialize in a specific task, such as decoupling perturbation response from cell state^4–6^, disentangling group-specific attributes^7^ or using disentangled representations for out-of-distribution sampling of single-cell data^8,9^. However, these are either task-specific and do not address the general disentanglement problem, rely on linearity and independence assumptions (and thus cannot capture nonlinear effects and interactions), cannot integrate multiple types of information beyond the single-cell measurements, or do not provide a generic reconstruction procedure.

**Figure 1:**
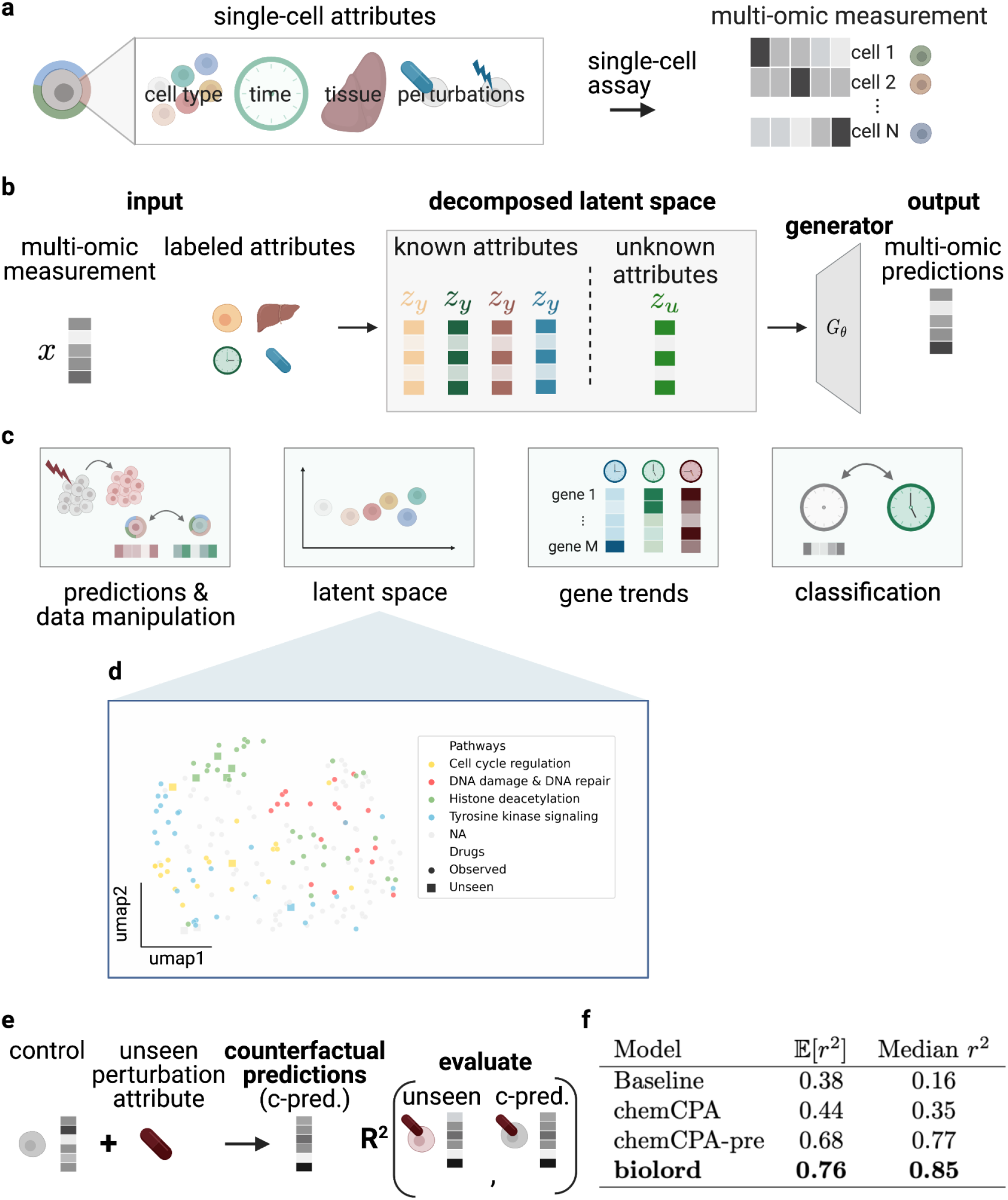
The biolord framework for disentanglement of known and unknown attributes. **(a)** Single-cell data encodes multiple attributes of cellular identity. **(b)** Schematic overview of the biolord model; Given single-cell measurements and labels for observed attributes, biolord finds a decomposed latent space and defines a generative model for measurement predictions. **(c)** Biolord can be used for multiple downstream tasks. From left to right: *predictions & data manipulation*; we can predict the gene expression of unseen cellular states, providing the unseen labels as input to the model coupled to a control cell, or similarly, study the changes in gene expression which correspond to a manipulation of a cell’s attribute. *latent space*; the decomposed latent space provides insight into the underlying structure of individual attributes. The independent latent space representation of each attribute reveals this structure. *Gene trends*; we can identify gene trends as a function of the known attributes. *classification*; using the semi-supervised biolord architecture we can label cells with missing attributes. **(d)** A UMAP^17^ of the learnt drug embedding on the highest dosage (10 *μM*). Dots, representing drugs, are colored according to known pathways. The shape represents whether the drug is *observed* (circle) or *unseen* (square). **(e)** Model evaluation procedure; control measurements are coupled with labels of unseen drugs to produce counterfactual predictions (c-pred.). *r*^2^ score is calculated between held-out samples and c-preds. **(f)** An evaluation of the model performance on predictions of unseen drugs on the highest dosage (10 *μM*), the strictest setting as it induces the most change in gene expression.

The disentanglement task is a cornerstone problem in machine learning, which has recently attracted much research interest^10–12^. This task aims to recover the underlying meaningful factors that shape a complex sample (e.g., a natural image). For example, given a dataset consisting of photos of cars, a disentanglement algorithm may be used to automatically recover the different attributes in each image; the model of the car, its color, viewing angles, and possibly, an additional embedding describing all remaining properties. As the task of disentangling the different attributes is impossible without any inductive bias^13^, a common approach is domain-supervised disentanglement. In such a setting, we have access to labels of some of the attributes we wish to disentangle, e.g. the model of the car and its color.

Here, we leverage recent advancements in the field of domain-supervised disentanglement from the computer vision field by extending deep learning frameworks that rely on latent optimization^14,15^. We present **biolord** (**biolo**gical **r**epresentation **d**isentanglement), a deep generative framework for learning disentangled representations of known and unknown attributes in single-cell data (Methods). To disentangle single-cell data into its underlying attributes, we assume partial supervision over known attributes along with single-cell measurements. For example, the known attributes may be cell type labels, measurement time, or perturbation values, and in general, may be categorical (discrete) or ordered (continuous). Given the partial supervision, biolord finds a decomposed latent space, encompassing informative embeddings for each known attribute and an embedding for the remaining unknown attributes in the data. Importantly, unlike the enhancement or filtering task, which provides a modification with respect to known attributes, here we obtain a decoupled representation for all (known and unknown) attributes in the data. The generative module can in turn use the decomposed latent space to predict single-cell measurements for different cell states across variations in internal or external conditions. The approaches we adapt^14,15^ learn a unique latent code to represent each known attribute and an additional latent code that describes all the unknown (unsupervised) attributes. The disentangled representation is obtained by inducing information constraints; the loss attempts to maximize the accuracy of the reconstruction (enforcing completeness) while minimizing the information encoded in the unknown attributes (limiting its capacity). Importantly, here we use an inductive bias accounting for the features of single-cell data (in contrast to images) through architecture and design choices and modify the framework for a set of biologically-meaningful downstream tasks (Fig. 1b-c, Methods).

The generality of the framework allows its application to diverse biological settings which can be studied with a rich set of downstream analysis tasks. The decomposed latent space allows studying the different attributes and their inner structure independently, for example, inferring similarities between drugs used for perturbations. Moreover, using the generative part of the model we can perform data manipulation, such as querying how a cell’s expression would have changed if it was sampled at a different time point or a different location in the tissue. Alternatively, we can make predictions for unobserved states, such as perturbation response to unseen drugs. More generally, the framework allows studying gene trends as a function of cell state (Fig. 1c, Methods). We implemented biolord using the scvi-tools library^16^ and made it available as open-source software at https://github.com/nitzanlab/biolord.

### Representation disentanglement accurately predicts cellular perturbation response

Accurate prediction of molecular responses to external perturbations is central to our understanding of cellular behavior and to translational medicine. Hence, many tools are developed dedicated to this task^5,6,18^. Amongst these is chemCPA^6^, an encoder-decoder neural network architecture that includes molecular information about drugs for response prediction. chemCPA resembles the biolord setting in terms of its input, however, is applicable specifically for drug perturbations tasks. The architecture is based on amortized training in contrast to the latent optimization scheme used by biolord which was shown to perform better^14,15^ (Methods).

Cellular response prediction can be framed as a disentanglement task, aimed at decoupling perturbation response from the underlying cell state, and therefore can be approached by biolord. We use the sci-Plex 3 dataset which includes ∼650,000 single-cell transcriptomes from 3 cancer cell lines exposed to 188 compounds at 4 different dosages and control samples^19^ (Supplementary Fig. 2). We consider a setting suggested by Hetzel et al.^6^, assessing the accuracy of predictions of perturbation response to nine unseen compounds (reported among the most effective drugs in sci-Plex 3 data^19^).

To allow biolord to generalize to unseen drugs, we take advantage of existing prior knowledge and obtain chemically informed embedding of the drugs using RDKit features^6,20^. For each cell, the features of each drug, alongside its dosage, the cell line, and corresponding scRNA-seq measurements, are given as input to biolord (Methods, Supplementary Note 1). First, we note that biolord’s learned latent representation is biologically informative and reveals drug organization according to known corresponding pathways (Fig. 1d). This representation better captures underlying drug organization, relative to the chemically informed RDKit features used as input, both qualitatively (Supplementary Figure 3) and quantitatively (Adjusted Rand Index *RDKit*: 0.03, *biolord*: 0.16). Next, this setting allows us to use the biolord model to obtain counterfactual predictions for unseen drugs. Namely, to generate the predicted expression of control cells with the labels of unseen compounds. To evaluate the performance of predicted perturbed states, we compute the *r*^2^ score between the real measurements of cells exposed to the unseen compounds and the counterfactual predictions (Fig. 1e, Methods). We consider the strictest setting, being prediction over the highest dosage (10 *μM*). Biolord outperforms both a naive baseline (comparing real measurements of unseen compounds to the control measurements), as well as state-of-the-art models, chemCPA and chemCPA-pre (a chemCPA model pre-trained using bulk RNA-seq high-throughput screens) (Fig. 1f; for performance on all dosage values see Supplementary Fig. 4). Importantly, biolord outperforms chemCPA-pre in predicting cellular response (the mean value for the highest dosage, 10 *μ M, chemCPA-pre*: 0.68 *biolord*: 0.76), though not provided with the additional information used by chemCPA for pre-training.

### Identifying gene programs modified by *Plasmodium* infection via counterfactual predictions

A trained biolord model can be used to make counterfactual predictions and generate gene expression patterns of cells with a modified attribute. For example, recovering changes in the expression of a given control cell had it been infected (Fig. 1c, Methods). Ideally, such counterfactual predictions should capture differences in gene expression induced only by the modified attribute, and thus reveal gene programs related to it. We showcase biolord’s ability to disentangle and obtain insightful counterfactual predictions in complex biological settings by next focusing on a spatio-temporal single-cell atlas of *Plasmodium* infection progression in the mouse liver^21^. Single-cell data, including host and parasite transcriptome, was collected from infected mice at five time points post-infection (2, 12, 24, 30, and 36 hours post-infection (hpi)), as well as from control mice, not exposed to the parasite (control) (Fig. 2a, Supplementary Fig. 5). To classify hepatocytes from injected mice as infected or uninfected, the authors relied on GFP content in the parasite transcriptome^21^ (Fig. 2b). Using biolord, we aim to decouple the changes in gene expression in the host hepatocytes caused by the infection process from the variability rooted in previously established spatiotemporal processes^22,23^: either in spatial zonation across liver lobules radial axis or in temporal variation along the time of day (Fig. 2a, Supplementary Fig. 6).

**Figure 2:**
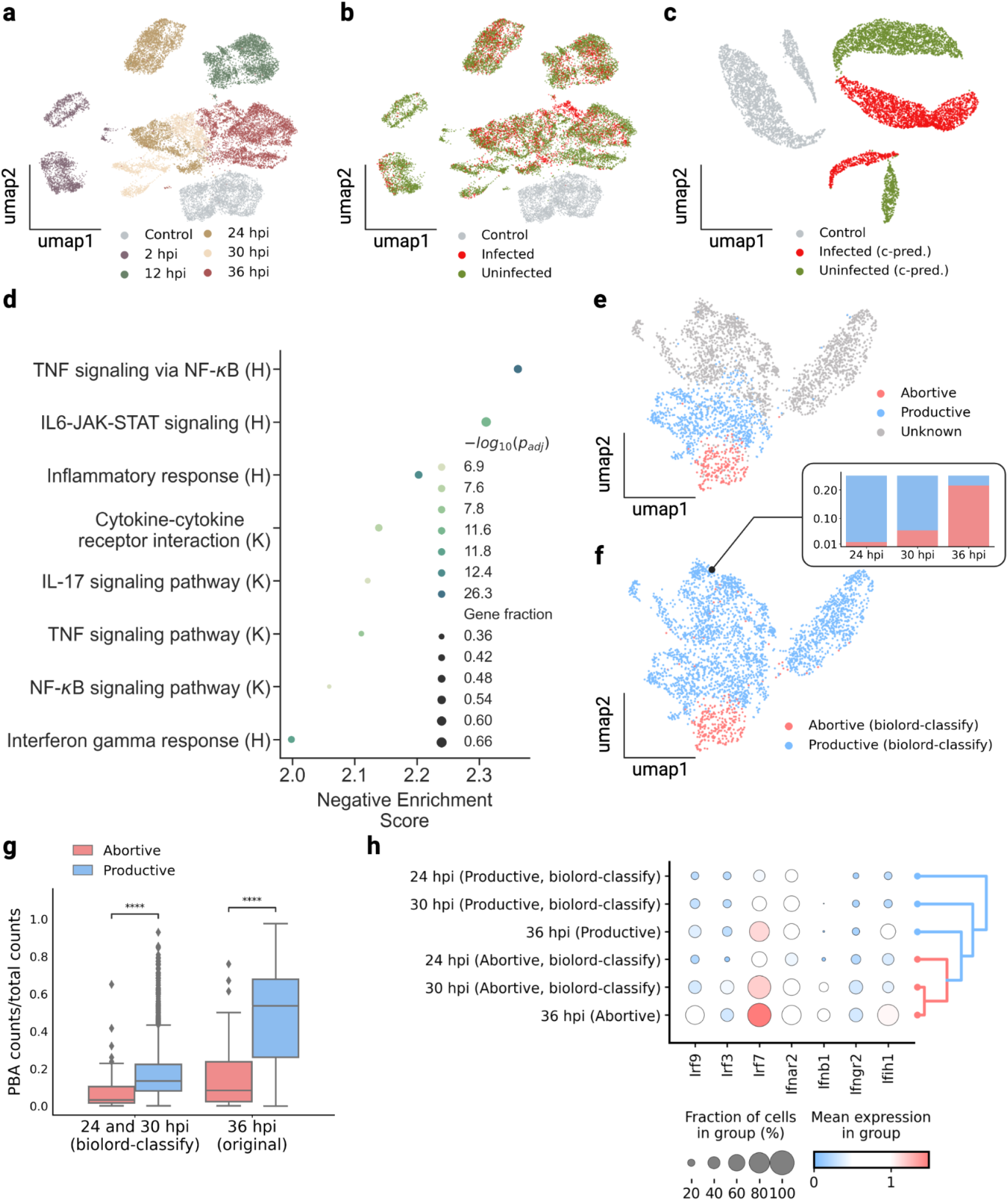
Recovering transient states by classifying unknown cell states using biolord. **(a)-(b)** UMAPs of the single-cell atlas of the *Plasmodium* liver stage^21^. Cells are colored by **(a)** time after infection, and **(b)** reported classification to infected/uninfected and control cells. **(c)** UMAP of the original control cells with their counterfactual predictions (c-pred.) for infected/uninfected state, cells are colored by the corresponding state. **(d)** Gene set enrichment analysis (GSEA) of genes found associated with the infected state based on biolord’s counterfactual predictions of the infection state in control cells. H, denotes Hallmark gene sets; K, KEGG gene sets. **(e)-(f)** UMAPs of the infected cells from intermediate to late time points in the single-cell atlas of the *Plasmodium* liver stage^21^. Cells are colored by, **(e)** the reported abortive/productive classification of cells at 36 hours-post-infection (hpi), and **(f)** biolord’s classification of all infected cells as abortive/productive. The inset shows the fraction of abortive cells at each time point (*24 hpi*: 0.016, *30 hpi*: 0.057, *36 hpi*: 0.215). **(g)** Boxplot comparing abortive and productive cells shows that abortive hepatocytes retain a smaller fraction of *Plasmodium* transcriptome across all time points (Mann-Whitney-Wilcoxon test two-sided with Benjamini-Hochberg correction: *24 and 30 hpi (biolord-classify)*: 8.192e-11, *36 hpi (original)*: 2.17e-53). **(h)** Abortive cells present an over-expression of interferon response as demonstrated by an increase in interferon regulatory factors and response-associated genes. The dendrogram ordering of the groups shows a trajectory from productive to abortive cells ordered by hpi.

To obtain counterfactual predictions for control cells with an infected status, we train a biolord model with infected and uninfected hepatocytes from injected mice, as well as hepatocytes from control mice. We provide as input the host transcriptome along with status classification (infected/uninfected/control), spatial zone (periportal/pericentral), and time (2, 12, 24, 30, 36 hpi or control) (Fig. 2c, Methods, Supplementary Note 2). These predictions, in turn, allow us to study directly how the infection state affects gene expression programs. To assess infection-related changes at the level of individual cells we use the dependent t-test for paired samples, testing for the null hypothesis that the counterfactual predictions and original expressions have identical average (expected) values. We performed the test for each gene and then used the results as input for gene set enrichment analysis (GSEA), which revealed an increase in the expression of genes associated with immune and stress pathways in infected hepatocytes (Fig. 2d). These findings are in accordance with previous reports^21^. However, in the original analysis, the comparisons between infected and uninfected hepatocytes had to be done for cells that were matched in terms of spatial lobule coordinates and sampling time, while biolord allows for global integrated analysis.

### Biolord exposes the transient trajectory towards distinct infection states

So far we have assumed full supervision over known attributes (meaning that for a known attribute, all cells are labeled), however, this is not always the case. Often only a subset of the cells is annotated. In such cases, we can leverage these partial labels to classify the remaining cells using biolord-classify, a biolord model coupled with a classifier for each attribute for the prediction of missing labels (Supplementary Fig. 1, Methods). The spatio-temporal single-cell atlas for *Plasmodium* infection presented above^21^ provides an example of such a setting. In their analysis, Afriat et al.^21^ identified a subpopulation of cells that shows a pattern of vacuole breakdown, termed ‘abortive hepatocytes‘. In the scRNA-seq data, this population was identified only at the latest time point (36 hpi) (Fig. 2e, Supplementary Fig. 7). However, in analyzing smFISH images, the existence of this population was verified as early as 24 hpi^21^. This provided the motivation to use biolord-classify to classify abortive cells within the scRNA-seq data at earlier time points, or in other words, identify the cells that would have progressed to become identifiably abortive at 36 hpi.

We train a biolord-classify model over hepatocytes at late time points (24, 30, and 36 hpi), where an abortive population is already expected to appear^21^. As input, we use the host transcriptome along with complete supervision over spatial zone (periportal/pericentral) and time (24, 30, 36 hpi), and partial state classification (abortive/productive/unknown) (Supplementary Note 2). The biolord-classify model allows us to label cells at earlier time points (24 and 30 hpi) as abortive or productive, thus predicting a temporally-extended abortive population (Fig. 2f, Methods). The extended abortive population preserves host gene expression trends observed in the original 36 hpi population, as expected. Namely, representative genes found to be upregulated in abortive hepatocytes at 36 hpi^21^ are statistically significantly upregulated in predicted abortive cells across all time points (Supplementary Fig. 8). Further, cells predicted to be abortive by biolord showcase reduced levels of *Plasmodium* transcripts and appear at earlier phases of *Plasmodium-*based pseudotime, consistent with findings regarding the original abortive population at 36 hpi^21^, even though these attributes were not used to train the model (Fig. 2g, Supplementary Fig. 9, Methods). Additionally, we recover the periportal bias of the abortive population (Supplementary Fig. 9).

The extended abortive population classified by biolord shows an increased IFN response across all time points, demonstrated by the over-expression of interferon regulatory factors (IRF3/7/9), which regulate the transcription of type I IFNs, and an increase in IFN*α* γ, genes (Fig. 2h). This is consistent with the hypothesis linking the abortive state to interferon-mediated innate immune response induced by *the Plasmodium* liver stage^21,24,25^. Further, biolord captures a transient trajectory of cellular states, showing a gradual increase in IFN signal across time within the abortive sub-population (Fig. 2h).

## Discussion

Here we presented biolord, a deep generative framework for learning disentangled representations of known and unknown attributes in single-cell data using partial supervision. We modify and extend established deep learning frameworks from the computer vision field to account for the inductive bias tailored for single-cell data and attend diverse biologically-meaningful analysis tasks. We have demonstrated biolord’s application to a wide variety of tasks showcasing the range of biological insights such disentangled representations can provide.

We started by applying biolord to the prediction of drug perturbation response and showed that while being a generic framework, it outperforms methods that are dedicated to this task which use additional prior knowledge, not provided to the biolord model. Next, by studying the spatio-temporal single-cell atlas of *Plasmodium* infection, we demonstrated biolord’s ability to decouple biological signals and recover gene expression trends. Further, for the case of partial supervision, we presented biolord-classify and demonstrated its ability to extend partial attribute supervision to the entire population, thereby recovering global features associated with the infection-related attribute and identifying transient states. Beyond the settings presented here, biolord can be applied to any multi-modal single-cell dataset for which partial supervision exists over a set of attributes. For example, a single-cell developmental atlas that includes samples from different tissues can be disentangled and manipulated, providing the age and tissue of origin as known attributes.

As with any deep generative framework, biolord suffers from the lack of direct interpretability. In addition, similarly to other disentanglement methods, it is unclear what is the desired outcome when known and unknown attributes are highly correlated. By providing a decomposed latent space, biolord allows extracting the underlying structure for each biological attribute independently, overcoming the above limitations. Further, the generative model exposes gene expression trends, providing interpretability in feature space. Thus, biolord provides a step towards decoupling cellular identities encoded in single-cell data; elucidating the effects of the different components on the overall observed expression, thereby providing new insights and better utilization of multi-omics data

## Methods

### Latent optimization as an inductive bias for disentanglement

Latent optimization is a critical component in our approach. Typical representation disentanglement approaches use an encoder to map the original data samples into latent codes. This is often called *amortized* inference. While having an encoder network to map samples to codes is convenient, Gabbay and Hoshen^14^ showed that this approach may achieve subpar results. The reason is that at the beginning of training, a randomly initialized encoder maps all sample attributes to each latent code, without any disentanglement. While the loss function encourages disentanglement, this poor initialization makes it hard to achieve disentangled results in practice, due to optimization challenges. On the other hand, randomly initialized latent codes, trivially do not contain any information on non-target attributes (or any other information). In the course of training, the latent codes become more informative over the target attribute, while the disentanglement objectives ensure that they do not gain information over other attributes. Intuitively, this is as preventing the gain of unwanted information is easier than losing existing information. To conclude, latent optimization helps achieve more disentangled latent codes by providing a better initialization for the learning process.

### Biolord: biological representation disentanglement

Consider a dataset 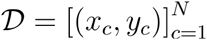 where for each cell *c, x* _*c*_ ∈ ℝ ^*M*^ describes the gene expression of *M* genes, and *y* _*c*_ is a set of known cell attributes such as cell-type, tissue of origin or age. We assume that the gene expression can be disentangled into representations in multiple latent spaces, each representing an observed cell attribute and a single space for unsupervised properties. As we elaborate later, within we make a distinction between categorical and ordered attributes when finding its latent space.

Broadly speaking, the biolord framework takes the dataset *D* and constructs a decomposed latent space using a set of dedicated neural networks. We denote *z*_*y*_ the output of each sub-network, the latent space representing each attribute (categorical or ordered) with *y* ∈ *γ*, and *z*_*u*_ the latent space of unknown attributes (Fig. 1b). The latent space embedding maps each sample to a latent code. It is important to note that we do not attempt to distinguish between the different unknown attributes. Next, a generator *G* is defined, taking as input the decomposed latent space and outputting predictions for the gene expression.

### Attributes’ latent space

For each attribute *y* ∈ *γ* we need to obtain *z*_*y*_, the latent space representing it. Here we make a distinction between categorical attributes, where similar cells share class labels, and the ordered attributes, in which distances between the attribute’s features encode similarity. Hence, for the categorical attribute, we model each attribute’s representation as an embedding *z*_*y*_, such that the latent code is shared between all cells belonging to the same label. We optimize these embeddings directly (namely, latent optimization) with the model objective. However, for the ordered attribute, we use encoders (an encoder for each attribute) - a multi-layer perceptron (MLP) that maps the features to the latent space *z*_*y*_.

### Latent space of unknown attributes using a per-sample embedding

We learn the unknown attributes’ representation by optimizing per-sample embeddings directly. That is, a latent code *z*_*u*_, is assigned to each cell, independent of gene expression or known attributes. This embedding is optimized during training using latent optimization. Yet, optimizing a unique code for each cell may hinder our disentanglement efforts: The model may encode the entire expression information with the latent code of unknown attributes, and ignore the attribute-specific encoding. To ensure that known attribute information does not leak into the representation of the unknown attribute, we regularize it in two manners. First, is inducing an additive Gaussian noise η ∼ Ν, for (0, *σ*^2^ *I*) some fixed variance value σ (‘unknown_noise_param‘). Secondly, we add an activation decay penalty, γ (‘unknown_penalty‘) to the loss, inducing the minimality loss term,

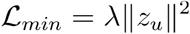

Together, these enforce the minimality of shared information between the representation of unknown attributes and known attributes. That is, the representation of unknown attributes is optimized to minimize the information it encompasses regarding known attributes.

### Expression generator

To reconstruct the underlying distribution of gene expression space from the decomposed latent space we define a generator, *G*. The generator is a neural network parameterized by *θ* which takes as input the concatenated decomposed latent space, and outputs a parametrization of the gene expression distribution (given by the mean and variance),

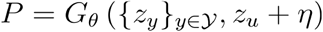

Depending on the data provided as input to the model, pre-processed or raw counts, the distribution,*P*, can follow a Gaussian distribution or zero-inflated negative binomial (ZINB), respectively^16^. To define the reconstruction, and completeness loss term, we use the respective negative log-likelihood loss for each distribution,*NLL*(*x*/*G*_*θ*_). For the Gaussian case, we add a mean squared error term, concerning the predicted means,*MSE*(*x*, μ*θ*), tuned by *τ* (‘reconstruction_loss‘) hyperparameter (set to 0 for the ZINB setting). Hence we can write the completeness term as

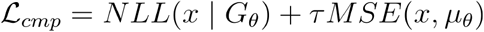

### Model objective

Combining the above we can write the model’s loss function as a composition of two terms. The first term induces completeness by optimizing the accuracy of the generator, and the second enforces the minimality of information shared between the representations of known and unknown attributes,

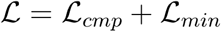

### Biolord-classify: biological representation disentanglement with partial labels

To perform semi-supervised disentangling, namely, a setting in which we have missing labels for a subset of cells, we adopt a setting presented by Gabbay et al.^15^ for semi-supervised image disentanglement. Based on the biolord model defined above, for each attribute, we train either a classifier (for categorical attributes) or a regressor (for ordered attributes), which takes as input the gene expression and outputs the class label/features. For cells with missing labels, we use the classifier’s (regressor’s) output to obtain the decomposed latent representation. Namely, the class latent space is used as input for the generator (Supplementary Fig. 1).

For each classifier (regressor), we add a term to encourage the correct prediction for the samples for which labels are available. For the classifiers, we use the categorical cross-entropy loss. For the regressors, we use the mean squared error loss between the output and provided features. In all cases, the loss is evaluated only over cells for which labels are provided.

### Datasets, training, and evaluation

#### sci-Plex 3

We consider the sci-Plex 3 data set presented in Srivatsan et al.^19^ which contains measurements for 649,340 cells across 7561 genes. The data contains three human cancer cell lines—A549, MCF7, and K562 with perturbations for 188 drugs at four different dosages 10 *nM*, 100 *nM*,1 *μM*, and 10 *μM*. We use a pre-processed anndata file provided by Hetzel et al.^6^, downloaded from https://f003.backblazeb2.com/file/chemCPA-datasets/sciplex_complete_middle_subset.h5ad. To the downloaded anndata file we add RDKit2d^20^ features using chemprop package^26^ and an out-of-distribution split, keeping nine unseen drugs for validation: Dacinostat, Givinostat, Belinostat, Hesperadin, Quisinostat, Alvespimycin, Tanespimycin, TAK-901, and Flavopiridol.

#### Training parameters

We train a biolord model over the processed gene expression (hence assuming Gaussian distribution). We use RDKit chemically informed features embedding of the drugs^27^, as well as the dosage as ordered attributes. The cell line is passed as a categorical attribute (detailed model configurations are given in Supplementary Note 1). We used Weights & Biases^28^ for experiment tracking and hyperparameter tuning (hyperparameters ranges and final hyperparameters reported are provided in Supplementary Note 1).

#### Evaluation and benchmarks

Following the procedure suggested by Hetzel et al.^6^ we evaluate the prediction accuracy using the coefficient of determination *r*^2^(*r*^2^score). The *r*^2^score is calculated between a model’s counterfactual predictions and the ground truth measurements on all genes.

For benchmarks, we consider three settings, a naive baseline, and two settings of the chemCPA framework^6^. For the naive baseline, we evaluate the *r*^2^score between control, unperturbed cells (per cell line), and the respective drug-treated cells. This provides a proxy for the complexity of the task, quantifying the effect of the perturbation on gene expression space. For chemCPA, we consider the standalone setting (chemCPA), which trains the drug encoding network directly on the single-cell data, and a pre-trained model, for which the drug encoding network was trained over bulk RNA high-throughput screen (L1000^29^) (chemCPA-pre). The pre-trained model was kindly shared with us by the authors of chemCPA^6^. To run the chemCPA benchmarks we rely on optimal hyperparameters reported in the original publication for the sci-Plex 3 setting. For chemCPA-pre we perform an additional round of hyperparameter tuning for all adversary parameters (hyperparameters ranges and hyperparameters used to report results for chemCPA and chemCPA-pre are provided in Supplementary Note 1).

#### spatio-temporal single-cell atlas of the *Plasmodium* liver stage

To study the liver stage of the malaria parasite *Plasmodium*, Afriat et al.^21^ molecularly characterized thousands of infected and uninfected hepatocytes. Infected cells were collected at five time points post-infection (2, 12, 24, 30, and 36). We downloaded the pre-processed annotated data provided by the authors from Zenodo^30^. The data annotations include:

- coarse_time: denoting the number of hours post-infection when the cells were collected (or control).
- eta_normalized: a spatial zonation score based on zonation marker genes which were used to classify the cells as periportal/pericentral.
- pseudotime: calculated using Monocle over the normalized data of the infected hepatocyte PBA genes subset.
- status: infection status inferred by FACS sorting of the hepatocytes.
- abortive: classification of cells at 36 hpi as Abortive/Productive based on clustering of host transcriptome.

#### Training parameters

We define two biolord settings, as described below.

#### Infected state analysis over the complete dataset

A biolord model is defined over infected and uninfected hepatocytes from injected mice, as well as hepatocytes from control mice (namely the dataset excluding Mock and Mosquito bitten samples). As input, we use the host transcriptome (restricted to 8355 genes used in the original publication) along with status classification (infected/uninfected/control), spatial zone (periportal/pericentral), and time (2, 12, 24, 30, 36 hpi or control). Hyperparameters for reported results are provided in Supplementary Note 2.

#### Abortive state classification

A biolord-classify model was trained over infected hepatocytes at 24, 30, and 36 hpi. As input, we use the host transcriptome (restricted to highly variable genes) along with spatial zone (periportal/pericentral), time (24, 30, and 36 hpi), a stress_score (computed using scanpy’s^31^ function ‘scanpy.tl.score_genes()‘ with stress genes reported by Afriat et al.^21^, defining an ordered attribute) and the partial abortive state classification given for 36 hpi (Abortive/Productive). Of note, we introduce the stress_score to disentangle the stress signal, reported in the original publication^21^, from the abortive signature. Hyperparameters for reported results are provided in Supplementary Note 2.

## Data availability

Datasets analyzed in this manuscript are publicly available. Processed data files can be downloaded from figshare (https://figshare.com/account/home#/projects/160085).

The original datasets analyzed in the current study are available at

- sci-Plex3:

https://f003.backblazeb2.com/file/chemCPA-datasets/sciplex_complete_middle_subset.h5ad, a pre-processed file provided by Hetzel et al.^6^.

- spatio-temporal single-cell atlas of the Plasmodium liver stage: publicly available at GSE181725 (https://www.ncbi.nlm.nih.gov/geo/query/acc.cgi?acc=GSE181725) or as processed Seurat object at https://zenodo.org/record/7081863.

## Code availability

We implemented biolord using the scvi-tools library^16^ and using cookiecutter-scverse (https://github.com/scverse/cookiecutter-scverse) as a template for the package. The package is released as open-source software at https://github.com/nitzanlab/biolord. Documentation is available at https://biolord.readthedocs.io. The code to reproduce the results is available at https://github.com/nitzanlab/biolord_reproducibility.

## Acknowledgments

We thank Amichay Afriat for his thoughtful review and assistance in analyzing the plasmodium liver stage single-cell atlas, Leon Hetzel for guidance in training the chemCPA models and for providing the pre-trained model for chemCPA, and Michal Klein for his thoughtful review and assistance in the software development. We acknowledge Bar Melinarskiy and all members of the Nitzan lab for general feedback.

