## Supplementary information for "Biological representation disentanglement of single-cell data"

Zoe Piran[1], Niv Cohen[1], Yedid Hoshen[1], and Mor Nitzan[1,2,3,*]

[1]School of Computer Science and Engineering, The Hebrew University of Jerusalem, Israel
[2]Racah Institute of Physics, The Hebrew University of Jerusalem, Israel
[3]Faculty of Medicine, The Hebrew University of Jerusalem, Israel
[*]

#### Contents

**Supplementary Figures** **2**

**Supplementary Note 1** **6**

sci-Plex 3 . . . . . . . . . . 6

**Supplementary Note 2** **10**

Spatio-temporal single-cell atlas of the Plasmodium liver stage . . . . . . . . . . 10

### Supplementary Figures

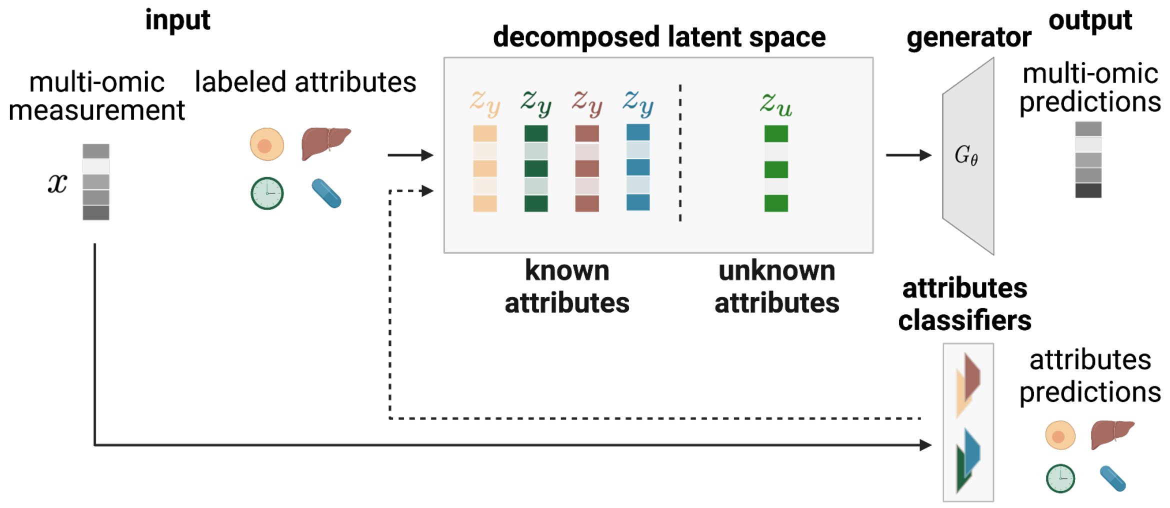


**Supplementary Figure 1: A schematic illustration of biolord-classify.** The semi-supervised biolord architecture. To handle partial labels we add classifiers to the naive biolord model (see Methods).

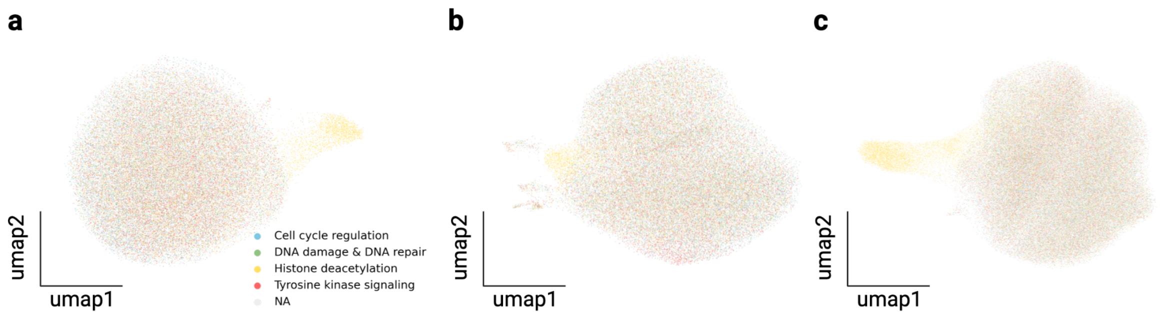


**Supplementary Figure 2: The sci-Plex 3 dataset[1].** **(a)-(c)** UMAPs of the original data seperated by cell-lines according **(a)** A549 (lung adenocarcinoma), **(b)** K562 (chronic myelogenous leukemia), and **(c)** MCF7 (mammary adenocarcinoma). Cells are colored by pathway associated with the drug-treated.

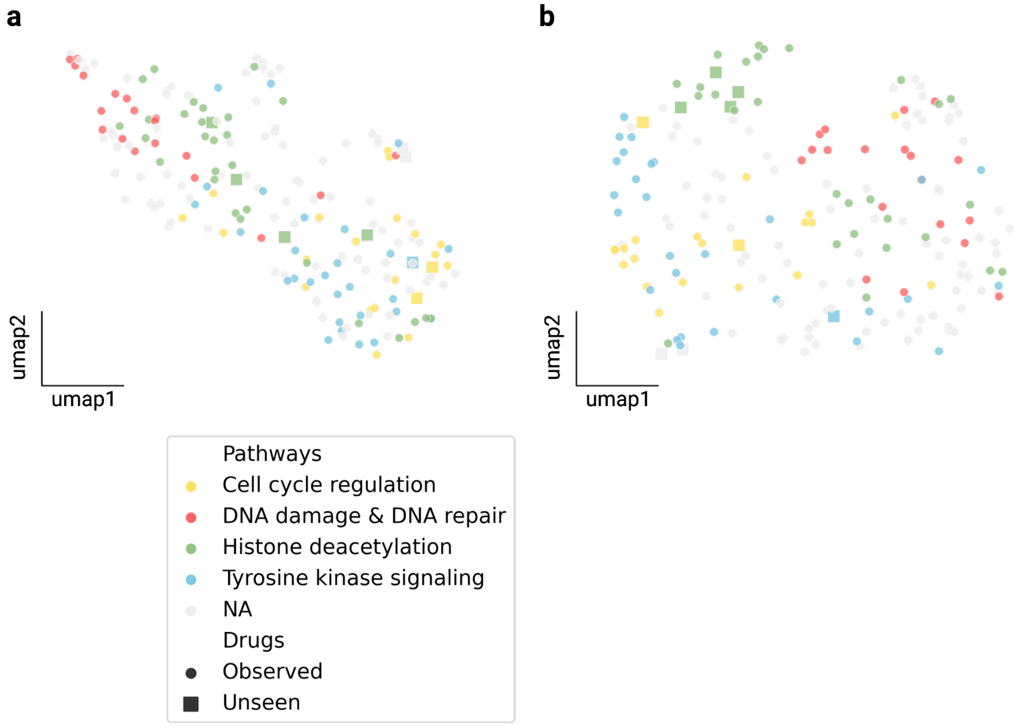


**Supplementary Figure 3: biolord decomposed latent space of the sci-Plex 3 dataset[1] exposes drug features.** **(a)** A UMAP of the chemically informed RDKit features used as input for biolord. **(b)** A UMAP of biolord's drug embedding on the highest dosage (10 $\mu M$). In both UMAPS, Dots, representing drugs, are colored according to known pathways. The shape represents whether the drug is *observed* (circle) or *unseen* (square).

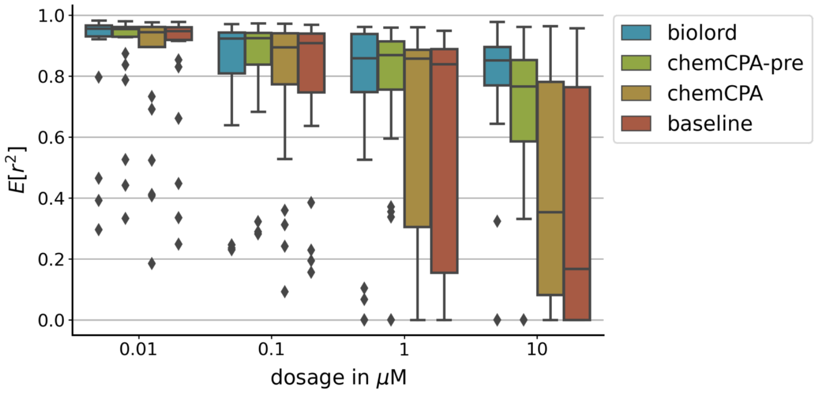


**Supplementary Figure 4: Benchmarking performance over the sci-Plex 3 dataset[1].** The mean $R^2$ score, over the nine unseen drugs and all genes. The score is reported for biolord, chemCPA pre-trained model (chemCPA-pre), the standard chemCPA (chemCPA), and the naive baseline (see Methods).

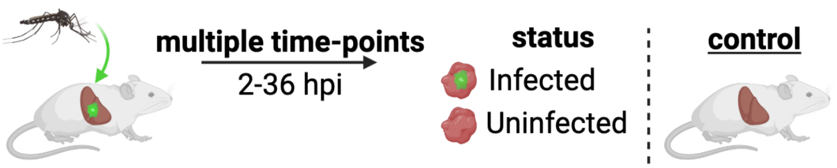


**Supplementary Figure 5: The single-cell atlas of the Plasmodium liver stage**[2]**.** Experimental schematic. GFP+ parasites are injected into mice and liver samples are extracted at different time points. Hepatocytes are classified as infected/uninfected using FACS sorting. Control samples are collected from healthy mice.

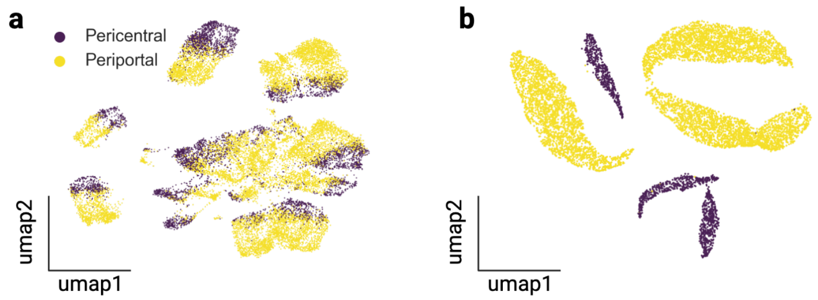


**Supplementary Figure 6: Zonation signature in counterfactual predictions of Plasmodium infection**[2]**.(a)** UMAP of the single-cell atlas of the Plasmodium liver stage; cells are colored by spatial zone. **(b)** UMAP of the original control cells with their counterfactual predictions (c-pred.) for infected/uninfected state; cells are colored by spatial zone.

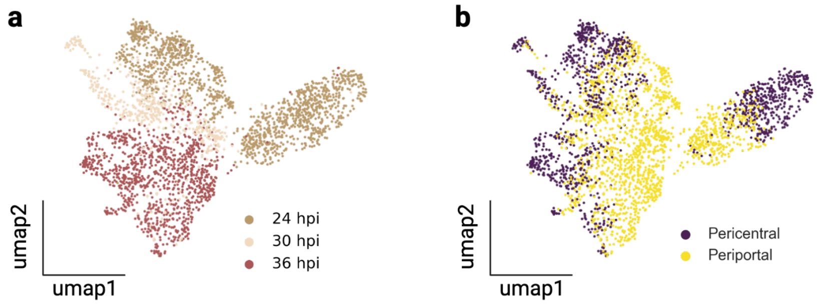


**Supplementary Figure 7: Application of biolord-classify to late time points in the Plasmodium liver stage atlas**[2]**.** **(a)-(b)** UMAP of cells from late time points; cells colored by **(a)** hours post-infection or **(b)** spatial zone.

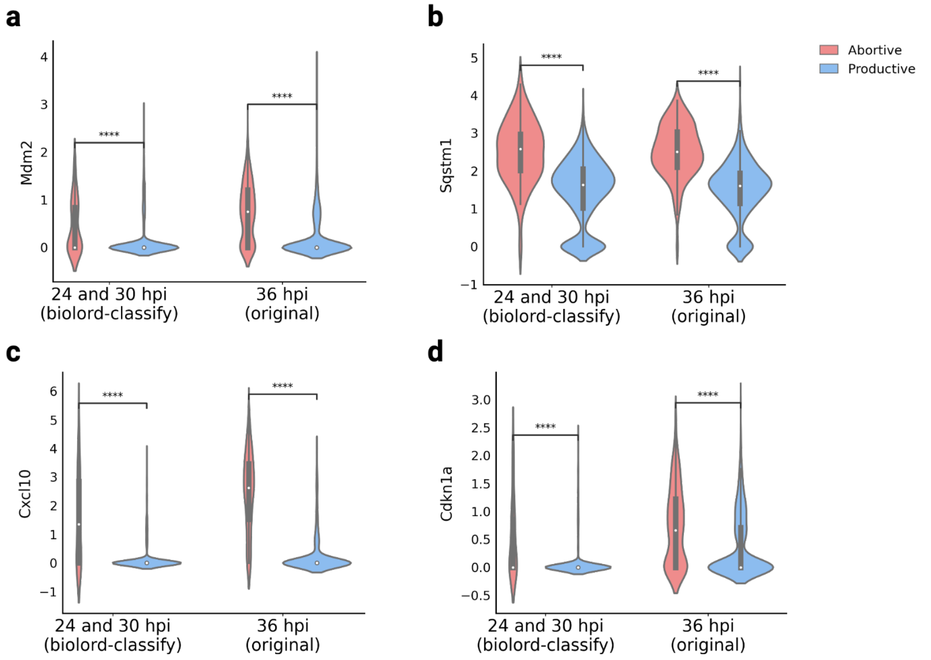


**Supplementary Figure 8: Gene expression patterns are recovered in abortive hepatocytes identified by biolord-classify**. Violin plots of representative genes upregulated in abortive hepatocytes. Mann-Whitney-Wilcoxon test two-sided with Benjamini-Hochberg correction P-values: **(a)** *Mdm2*; 24 and 30 hpi (biolord-classify): 2.786e-12, 36 hpi (original): 8.544e-47, **(b)** *Sqstm1*; 24 and 30 hpi (biolord-classify): 1.035e-14, 36 hpi (original): 1.27e-63, **(c)** *Cxcl10*; 24 and 30 hpi (biolord-classify): 7.201e-36, 36 hpi (original): 1.557e-117,**(d)** *Cdkn1a*; 24 and 30 hpi (biolord-classify): 3.71e-28, 36 hpi (original): 3.04e-17.

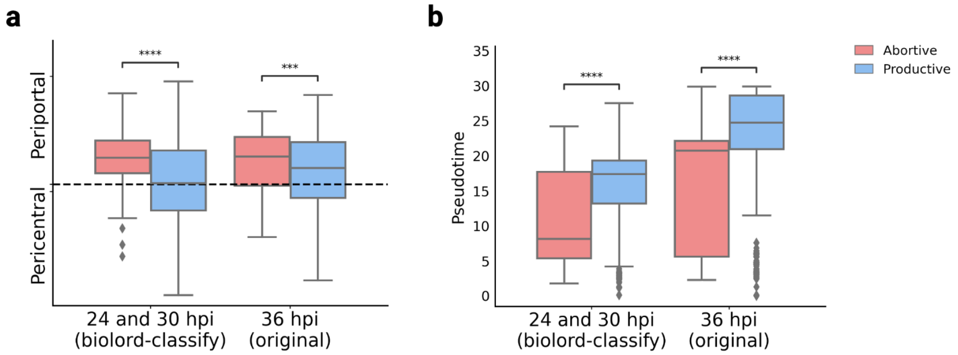


**Supplementary Figure 9: Global features are preserved in abortive hepatocytes identified by biolord-classify**. Boxplots comparing abortive and productive cells show that **(a)** abortive hepatocytes are more periportally zonated compared with productive hepatocytes; the $y$ axis represents zonation score and scores corresponding to Periportal/Pericentral spatial zones are indicated (see Methods, Mann-Whitney-Wilcoxon test two-sided with Benjamini-Hochberg correction P-values: *24 and 30 hpi (biolord-classify)*: 6.546e-05, *36 hpi (original)*: 3.056e-04). **(b)** The abortive population is concentrated at early pseudotime. Pseudotime was evaluated over parasite mRNA (Afriat et al. 2022) (Mann-Whitney-Wilcoxon test two-sided with Benjamini-Hochberg correction P-values: *24 and 30 hpi (biolord-classify)*: 1.165e-07, *36 hpi (original)*: 2.715e-35).

### Supplementary Note 1

#### sci-Plex 3

**biolord**. For all biolord models of the sci-Plex 3 dataset[1] we use the same anndata file available on figshare (sciplex3) and the biolord model settings as listed in Table 1. We used Weights & Biases[3] for experiment tracking and hyper-parameter tuning. To cover the large space of possible configuration space we tuned subsets of the parameters in consecutive iterations (Table 2). We started with tuning the more dominant parameters. For example, "loss_ae" or architecture parameters such as "n_latent", and "{}_depth" or "{}_width" parameters. We next tuned the loss parameters (e.g., penalties strength) and the dropout rate. Finally, we tuned the finer optimization parameters such as the learning rates and regularization parameters. We randomized over valid values for the inspected parameters at each iteration and kept the ones that consistently outperformed. The range of values scanned for each parameter and the best configurations are given in Table 2.

**Table 1:** General settings for a biolord model on the sci-Plex 3 data[1].

| Parameter | Value |
|---|---|
| ordered_attributes_keys | ["rdkit2d_dose"] |
| discrete_attributes_keys | ["cell_type"] |
| train_classifiers | False |
| batch_size | 512 |
| split_key | "split_ood" |
| max_epochs | 500 |
| early_stopping | True |
| loss_ae | "gauss" |

**chemCPA.** For the non-pre-trained version, we follow the parameters supplied in 'finetuning_num_genes.json' ('_id = 1007')[4]. The parameters are presented in Table 3.

**ChemCPA-pre.** For the pre-trained version, we followed 'finetuning_num_genes.json'[4] ('_id = 789'). As advised by the authors we tune the adversarial parameters, as detailed in Table 4.

**Table 2:** Parameters range for the biolord sweep on the sci-Plex 3 data[1] and optimal configuration used.

| Parameter | Values | Best config |
|---|---|---|
| n_latent | [16,32,64,128] | 256 |
| n_latent_attribute_ordered | [128,256,512] | 256 |
| n_latent_attribute_discrete | [2,3,4,6,8] | 3 |
| autoencoder_width | [32,64,128,256,512,1024,2048,4096] | 4096 |
| autoencoder_depth | [1,2,3,4,6,8] | 4 |
| autoencoder_lr | $[1 \times 10^{-2}, 1 \times 10^{-3}, 1 \times 10^{-4}]$ | $1 \times 10^{-4}$ |
| autoencoder_wd | $[1 \times 10^{-2}, 1 \times 10^{-3}, 1 \times 10^{-4}]$ | $1 \times 10^{-4}$ |
| attribute_nn_width | [32,64,128,256,512,1024,2048,4096] | 2048 |
| attribute_nn_depth | [1,2,3,4,6,8] | 2 |
| attribute_nn_lr | $[1 \times 10^{-2}, 1 \times 10^{-3}, 1 \times 10^{-4}]$ | $1 \times 10^{-2}$ |
| attribute_nn_wd | $[1 \times 10^{-8}, 4 \times 10^{-8}, 1 \times 10^{-7}]$ | $4 \times 10^{-8}$ |
| unknown_attribute_noise_param | $[0.1, 0.5, 1, 2, 5, 10, 20]$ | 20 |
| reconstruction_penalty | $[1 \times 10^{-2}, 1 \times 10^{-1}, 1 \times 10^{0}, 1, 1 \times 10^{2}, 5 \times 10^{3}, 1 \times 10^{4}]$ | $1 \times 10^{4}$ |
| unknown_attribute_penalty | $[0.1, 1, 2, 5, 10, 20, 50, 100, 200]$ | 0.1 |
| step_size_lr | $[45, 90, 180]$ | 45 |
| use_batch_norm | [True, False] | False |
| use_layer_norm | [True, False] | False |
| cosine_scheduler | [True, False] | True |
| attribute_dropout_rate | [0.05,0.1,0.25,0.5,0.75] | 0.1 |
| scheduler_final_lr | $[1 \times 10^{-3}, 1 \times 10^{-4}, 1 \times 10^{-5}, 1 \times 10^{-6},]$ | $1 \times 10^{-5}$ |

**Table 3:** Hyperparameters used for reported chemCPA results on the sci-Plex 3 data[1].

| Parameter | Value |
|---|---|
| num epochs | 200 |
| patience | 50 |
| dim | 32 |
| dropout | $2.624 \times 10^{-1}$ |
| autoencoder_width | 256 |
| autoencoder_depth | 4 |
| autoencoder_lr | $1.575 \times 10^{-3}$ |
| autoencoder_wd | $6.251 \times 10^{-7}$ |
| adversary_width | 128 |
| adversary_depth | 3 |
| adversary_lr | $8.060 \times 10^{-4}$ |
| adversary_wd | $4.000 \times 10^{-6}$ |
| reg_adversary | $9.101 \times 10^{0}$ |
| reg_adversary_cov | $1.068 \times 10^{1}$ |
| penalty_adversary | $4.550 \times 10^{-1}$ |
| batch_size | 32 |
| dosers_width | 64 |
| dosers_depth | 3 |
| dosers_lr | $1.575 \times 10^{-3}$ |
| dosers_wd | $6.251 \times 10^{-7}$ |
| embedding_encoder_width | 128 |
| embedding_encoder_depth | 4 |
| append_ae_layer | True |
| enable_cpa_mode | False |
| reg_multi_task | 0 |

**Table 4:** Parameters range for the chemCPA-pre sweep on the sci-Plex 3 data[1] and optimal configuration used.

| Parameter | Values | Best config |
|---|---|---:|
| num epochs | - | 200 |
| patience | - | 50 |
| dim | - | 32 |
| dropout | - | $2.624 \times 10^{-1}$ |
| autoencoder_width | - | 256 |
| autoencoder_depth | - | 4 |
| autoencoder_lr | - | $2.051 \times 10^{-4}$ |
| autoencoder_wd | - | $2.940 \times 10^{-8}$ |
| reg_adversary_cov | - | 4.176 |
| batch_size | - | 32 |
| dosers_width | - | 64 |
| dosers_depth | - | 3 |
| dosers_lr | - | $2.051 \times 10^{-4}$ |
| dosers_wd | - | $2.940 \times 10^{-8}$ |
| embedding_encoder_width | - | 128 |
| embedding_encoder_depth | - | 4 |
| append_ae_layer | - | True |
| enable_cpa_mode | - | False |
| reg_multi_task | - | 0 |
| adversary_width | $[64, 128, 256]$ | 256 |
| adversary_depth | $[2, 3, 4]$ | 3 |
| adversary_lr | $(5 \times 10^{-5}, 1 \times 10^{-2})$ | $1.143 \times 10^{-4}$ |
| adversary_wd | $(1 \times 10^{-8}, 1 \times 10^{-2})$ | $4 \times 10^{-6}$ |
| adversary_steps | $[2, 3]$ | 2 |
| reg_adversary | $(5, 100)$ | 1.778 |
| penalty_adversary | $(0.5, 5)$ | $8.89 \times 10^{-2}$ |

### Supplementary Note 2

#### Spatio-temporal single-cell atlas of the Plasmodium liver stage

As described in the main text we define two biolord settings for the analysis of the spatio-temporal single-cell atlas of the Plasmodium liver stage[2]. Below we provide the model details for each case.

##### Infected state analysis using counterfactual predictions

The relevant anndata file can be downloaded from figshare (spatio-temporal-infection_infected). The setting for the biolord model is provided in Table 5 and hyperparameter choice in Table 6. Of note, parameters relating to ordered attributes are missing as we do not have ordered attributes.

**Table 5:** The setting for the biolord model for the infected state analysis.

| Parameter | Value |
|---|---|
| discrete_attributes_keys | ["time_int", "status_control", "zone"] |
| train_classifiers | False |
| batch_size | 512 |
| split_key | "split_random" |
| max_epochs | 500 |
| early_stopping | True |
| early_stopping_patience | 20 |

**Table 6:** Parameters of the biolord model used for the infected state analysis.

| Parameter | Value |
|---|---|
| n_latent | 32 |
| n_latent_attribute_discrete | 4 |
| autoencoder_width | 1024 |
| autoencoder_depth | 4 |
| autoencoder_lr | $1 \times 10^{-4}$ |
| autoencoder_wd | $1 \times 10^{-4}$ |
| unknown_attribute_noise_param | $1 \times 10^{-1}$ |
| reconstruction_penalty | $1 \times 10^{2}$ |
| unknown_attribute_penalty | $1 \times 10^{1}$ |
| step_size_lr | 45 |
| use_batch_norm | False |
| use_layer_norm | False |
| cosine_scheduler | True |
| attribute_dropout_rate | 0.1 |
| scheduler_final_lr | $1 \times 10^{-5}$ |
| loss_ae | "gauss" |

##### Abortive state classification

The relevant anndata file can be downloaded from figshare (spatio-temporal-infection_abortive), named 'spatio-temporal-infection_abortive'. The setting for the biolord model is provided in Table 7 and hyperparameter choice in Table 8.

**Table 7:** The setting for the biolord model for the abortive state analysis.

| Parameter | Value |
|---|---|
| ordered_attributes_keys | ["stress_score"] |
| discrete_attributes_keys | ["time_int", "abortive_state", "zone"] |
| categorical_attributes_missing | {"time_int": None, "abortive_state": "Unknown", "zone": None} |
| train_classifiers | True |
| batch_size | 256 |
| split_key | "split_random" |
| max_epochs | 500 |
| early_stopping | True |
| early_stopping_patience | 20 |

**Table 8:** Parameters of the biolord model used for the infected state analysis.

| Parameter | Value |
|---|---:|
| n_latent | 32 |
| n_latent_attribute_discrete | 4 |
| n_latent_attribute_ordered | 16 |
| autoencoder_width | 512 |
| autoencoder_depth | 4 |
| autoencoder_lr | $1 \times 10^{-4}$ |
| autoencoder_wd | $1 \times 10^{-4}$ |
| attribute_nn_width | 512 |
| attribute_nn_depth | 4 |
| attribute_nn_lr | $1 \times 10^{-2}$ |
| attribute_nn_wd | $4 \times 10^{-8}$ |
| unknown_attribute_noise_param | $1 \times 10^{-1}$ |
| reconstruction_penalty | $1 \times 10^{2}$ |
| unknown_attribute_penalty | $1 \times 10^{1}$ |
| step_size_lr | 90 |
| use_batch_norm | False |
| use_layer_norm | False |
| cosine_scheduler | True |
| attribute_dropout_rate | 0.05 |
| scheduler_final_lr | $1 \times 10^{-5}$ |
| loss_ae | "gauss" |
| loss_ordered_attribute | "gauss" |
| classification_penalty | 0 |
| classifier_dropout_rate | $1 \times 10^{-1}$ |
| classifier_penalty | $1 \times 10^{1}$ |
| classify_all | False |
